# Uncovering differential tolerance to deletions versus substitutions with a protein language model

**DOI:** 10.1101/2024.06.27.601077

**Authors:** Grant Goldman, Prathamesh Chati, Vasilis Ntranos

**Affiliations:** Biological and Medical Informatics Program, University of California, San Francisco; Diabetes Center, University of California, San Francisco; Bakar Computational Health Sciences Institute, University of California, San Francisco; Department of Epidemiology & Biostatistics, University of California, San Francisco; Department of Bioengineering & Therapeutic Sciences, University of California, San Francisco

**Keywords:** Deletion scanning, protein language model, secondary structure, sequence context

## Abstract

Deep mutational scanning (DMS) experiments have been successfully leveraged to understand genotype to phenotype mapping, with broad implications for protein engineering, human genetics, drug development, and beyond. To date, however, the overwhelming majority of DMS have focused on amino acid substitutions, excluding other classes of variation such as deletions or insertions. As a consequence, it remains unclear how indels differentially shape the fitness landscape relative to substitutions. In order to further our understanding of the relationship between substitutions and deletions, we leveraged a protein language model to analyze every single amino acid deletion in the human proteome. We discovered hundreds of thousands of sites that display opposing behavior for deletions versus substitutions, i.e. sites that can tolerate being substituted but not deleted, and vice versa. We identified secondary structural elements and sequence context to be important mediators of differential tolerability at these sites. Our results underscore the value of deletion-substitution comparisons at the genome-wide scale, provide novel insights into how substitutions could systematically differ from deletions, and showcase the power of protein language models to generate biological hypotheses *in-silico*. All deletion-substitution comparisons can be explored and downloaded at https://huggingface.co/spaces/ntranoslab/diff-tol.

## Introduction

Understanding the effects of non-synonymous mutations is a critical challenge in fields such as human genetics and protein engineering^1,2^. To date, a growing computational and experimental literature has assessed the consequences of amino acid substitutions among a wide array of proteins^3–6^, yielding fundamental insights into how missense mutations are tolerated within a sequence context. However, other types of mutations, such as in-frame deletions, are far less understood^7,8^. Growing evidence suggests that deletions play a critical role in protein evolution and disease^8,9^, but it is unclear how a site’s substitution tolerance may differ from its deletion tolerance, if at all.

Although notable progress has been made in expanding experimental frameworks to incorporate deletions,^7,10^ they still remain vastly understudied relative to substitutions. According to a comprehensive survey of existing DMS data^6^, there are approximately seven thousand in-frame deletions, compared to over two million substitutions measured in the literature. Thus, current experimental data provides only a narrow window into the deletion fitness landscape.

Computational variant effect predictors (VEP) pose an attractive supplement to sparse experimental data on deletions. In particular, protein language models have recently emerged as efficient, scalable, and accurate predictors of mutation effects.^3,11–14^ Here we propose using a protein language model (pLM) as a proxy of experimental deletion scans, allowing us to comprehensively assess their genome-wide effects. We argue that, while individual pLM predictions may be noisy, collectively, at the genome-wide scale, they can generate a more representative view of the protein fitness landscape than limited experimental data alone. To this end, we employed ESM-1b^15^ to score every possible single amino acid deletion across all 42,161 protein sequences in the UniProt database^16^, resulting in a dataset of over 24 million unique deletion effects.

To showcase the use of protein language models as *in-silico* hypothesis generation tools, we focus on the property of differential tolerance – whether a site can tolerate a substitution but not a deletion, or vice versa (**Fig. 1**). Using our ESM-generated data, we identified 386,000 sites that meet this definition. We find that structural elements and sequence homogeneity are strongly associated with sites that are differentially tolerant. We further find evidence of differential tolerance in existing DMS studies and population genetics datasets, underscoring the importance of considering both deletions and substitutions in studies of protein fitness.

**Figure 1:**
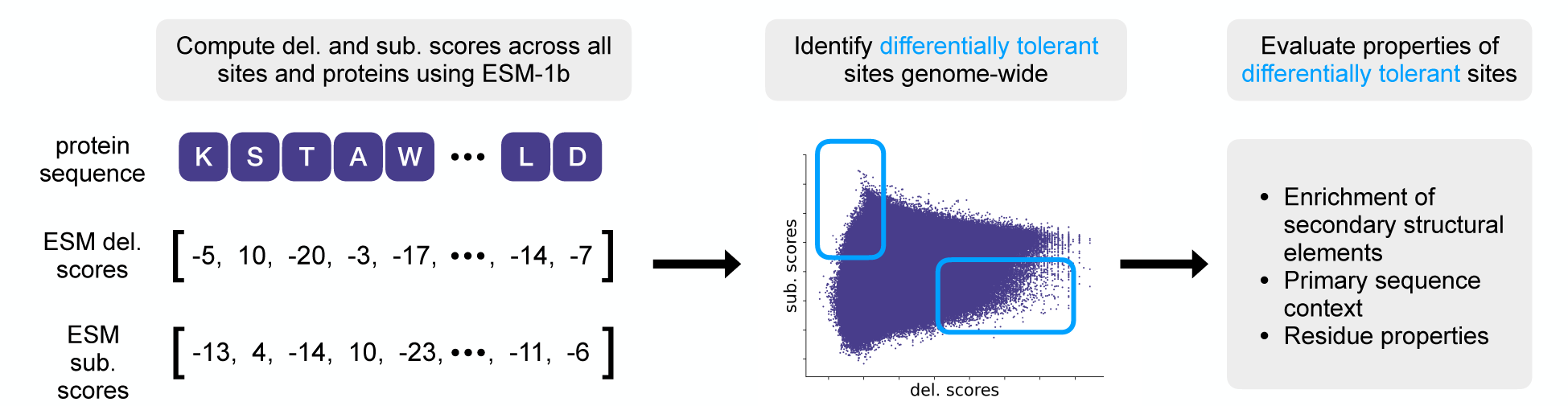
Genome-wide comparison of deletions versus substitutions with ESM. Overview of data generation process with ESM-1b, identification of sites with differential tolerance, and downstream analysis.

## Results

### Systematic comparison of substitutions versus deletions identifies sites with differential tolerance

We used our in-silico deletion dataset to estimate every site’s simultaneous tolerance to deletions and substitutions (as measured with ESM likelihood ratio scores, **methods**). For an individual site, while deletions and substitutions are generally correlated, our deletion scan identified many notable exceptions (**Fig. 2a**). In particular, we partitioned the substitution and deletion scores into their respective top and bottom quartiles and found 386,000 sites that rank among the 25% most damaging deletions *and* the 25% most neutral substitutions (red quadrant, **Fig. 2a**), or vice versa (green quadrant, **Fig. 2a**). We define the red quadrant as sites that are deletion-intolerant and substitution-tolerant, and refer to them as D^—^S^+^. Similarly, we define the green quadrant as sites that are deletion-tolerant and substitution-intolerant, and refer to them as D^+^S^—^.

**Figure 2:**
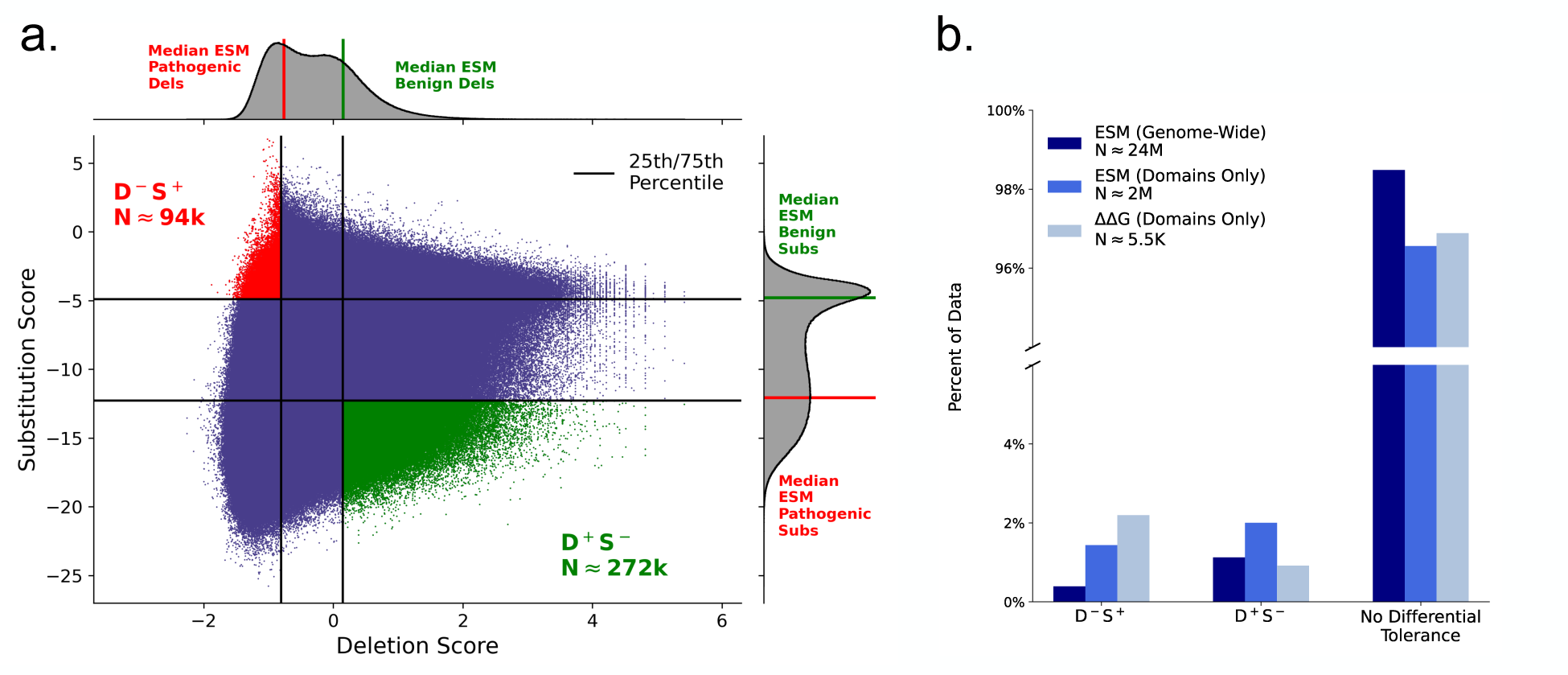
Identification of sites with differential tolerance. **(a)** Scatterplot of all site’s deletion versus substitution tolerance, sites with differential tolerance highlighted. Marginal distribution of genome-wide deletion scores (gray, top x-axis), with median ESM score of ClinVar benign deletions and pathogenic deletions highlighted. Analogous data for substitutions on the right y-axis. **(b)** Percent of sites with differential tolerance in ESM data (genome-wide and domains only) and experimental data (domains only)^10^.

Using ClinVar^17^ annotations we observe that every deletion score in the D^—^S^+^ quadrant is predicted more damaging than the median score among ClinVar pathogenic deletions, while substitution scores of these sites are predicted more neutral than the median score among ClinVar benign substitutions. Analogous results hold for the D^+^S^—^ quadrant (**methods, Fig. 2a**).

Notably, the proportion of ESM-predicted D^—^S^+^ sites and D^+^S^—^sites are consistent with estimates from experimental data (**Fig. 2b**). In particular, we analyzed changes in stability measurements (ΔΔG) across 177 unique functional domains (5,460 total sites) for both deletions and substitutions^10^. We partitioned the experimental data into their respective top and bottom quartiles and found 2.2% of sites meet the D^—^S^+^ criteria, and 0.92% of sites meet the D^+^S^—^criteria. These proportions are of similar order compared to the ESM-generated estimates of 0.4% and 1.1% in the genome-wide dataset, and 1.4% and 2.0% among sites in UniProt-annotated functional domains (**Fig. 2b**).

### Secondary structural elements mediate D^—^S^+^ differential tolerance

To understand what drives differential tolerance, we compared the frequency of secondary structural elements within the D^—^S^+^ and D^+^S^—^ quadrants against their background, genome-wide frequencies^16,18,19^. We first focused on the D^—^S^+^ quadrant and observed an enrichment of sites contained within alpha-helices (34.8% of sites genome-wide versus 62.2% among D^—^S^+^ sites, **Fig. 3a** and **Supplementary Fig. S1**). We reasoned this is a consequence of the periodic nature of the structure, which completes a full turn every 3.6 residues; substitutions conserve the number of residues in the turn, but deletions do not.

**Figure 3:**
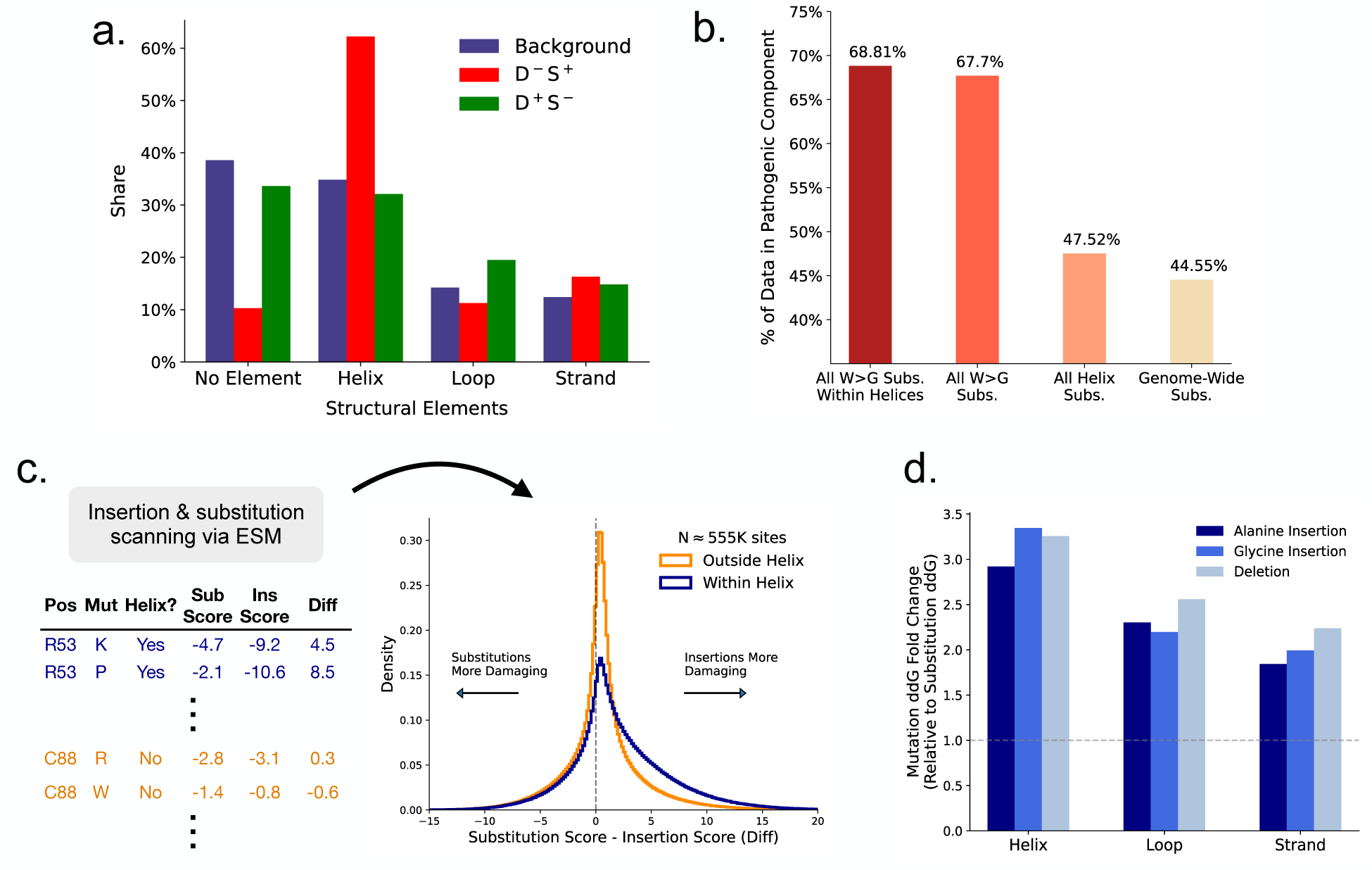
Secondary structural elements mediate differential tolerance to deletions versus substitutions. **(a)** The distribution of secondary structural elements genome-wide (blue), among D^—^S^+^ sites (red), and among D^+^S^—^ sites (green). **(b)** Comparison of W>G substitution score within helices versus all substitutions within helices, all W>G substitutions (inside and outside helices) *and* genome-wide effects. Each dataset was decomposed into a two-component Gaussian mixture, and the size of the inferred pathogenic component is shown. **(c)** Comparison of substitution effects versus insertion effects within and outside helices across 1,632 proteins. **(d)** Average change in experimentally-measured stability from insertions and deletions, normalized by average change in stability from substitutions, within secondary structural elements^10^.

Indeed, past work has investigated the effects of deletions within alpha-helices^10,20–23^, noting increased sensitivity to deletions. However, our analysis is the first to identify differential tolerance between substitutions and deletions within the structure. Jointly considering deletions and substitutions further clarifies the effect of deletions on alpha-helices. (**Supplementary Fig. S2).**

We also explored the model’s internal consistency with respect to D^—^S^+^ predictions by analyzing tryptophan (the largest amino acid) to glycine (the smallest amino acid) substitutions (W>G). This substitution, much like a deletion, removes mass and volume from the protein due to the difference in size between the amino acids. Because ESM-1b predicts alpha-helices to harbor the majority of D^—^S^+^ sites, we hypothesized that W>G substitutions should behave similarly to deletions within the structure. To quantify the effect of this substitution, we decomposed the distribution of scores into a two-component mixture of Gaussians, with one component representing an (inferred) pathogenic effect and the other an (inferred) benign component (**Supplementary Fig. S3**). We observe an increase in predicted pathogenic W>G substitutions within helices compared to its genome-wide effect (**Fig. 3b**). This result is notable as the substitution is likely enriched for damaging effects at baseline due to the dissimilarity of the amino acids^24–26^. Moreover, we demonstrate this increase in deleteriousness is not due to the mere presence of an alpha helix, as the pathogenic component of W>G substitutions within helices is larger than the pathogenic component of substitution across all pairs of residues within a helix, and the background effect of substitutions genome-wide (**Fig. 3b**). Overall, we identify similar directional effects between deletions and W>G substitutions within helices, suggesting ESM-1b has learned alpha-helices’ sensitivity to size-altering mutations.

To further probe the sensitivity of helices to length-altering mutations, we compared the ESM-predicted effects of insertions versus substitutions across 1,632 structurally diverse proteins **(methods)**. Relative to the effects of substitutions, insertions tend to be significantly more damaging within helices than outside helices (p<0.001, **Fig. 3c)**. Similarly, while insertions, deletions, and substitutions all tend to be more damaging within helices, this property is far more pronounced for insertions and deletions, which are length-altering, compared to substitutions (**Supplementary Fig. S4**).

As a final validation, we analyzed experimental stability measurements^10^ and found that insertions and deletions within helices are approximately three times as destabilizing as substitutions within helices. Helices show the clearest signal of differential tolerance to insertions and deletions (relative to substitutions) among all structural elements considered (**Fig. 3d**).

Taken together, these results clarify the tendency of alpha-helices to be overrepresented among sites that are simultaneously deletion-intolerant and substitution-tolerant. Our analyses indicate that ESM-1b has learned a critical underlying property of alpha-helices, namely that their periodicity renders size-altering mutations more deleterious compared to size-conservative mutations.

### Sequence context mediates D^+^S^—^ differential tolerance

Unlike sites that are deletion-*intolerant* and substitution-*tolerant* (D^—^S^+^, red quadrant, **Fig. 2b**), sites that tolerate deletions but not substitutions (D^+^S^—^, green quadrant, **Fig. 2b**) do not segregate as unambiguously by secondary structure (**Fig. 3a**). In order to understand why certain sites are deletion-tolerant and substitution-intolerant, we identified sequences enriched for this property. Among our examples, we observed many regions composed of chemically similar (or identical) amino acids. For example, the BRD2 protein (UniProt accession P25440) contains a 30-residue region, with 12 sites meeting the D^+^S^—^ criteria. The amino acids in this region are small and polar, with 21 serines and four aspartic acids. We therefore considered whether D^+^S^—^ sites were more likely to emerge within homogenous sequence contexts; as in the above example, one deletion from a string of many small, polar residues might be less damaging than substituting in a large, hydrophobic residue.

To test this, we needed to quantify the homogeneity of a given sequence. We first used BLOSUM substitution scores, which can be used to approximate a similarity measure between neighboring residues. Under this definition of homogeneity, we find that D^+^S^—^ sites are more similar to their neighboring residues than both D^—^S^+^ sites and random positions sampled across all proteins (**Fig. 4a**, **Supplementary Figs. S6** and **S7**). As in our secondary structure analysis, we find that this property is clarified by the joint analysis of deletions alongside substitutions. (**Supplementary Fig. S8)**.

**Figure 4:**
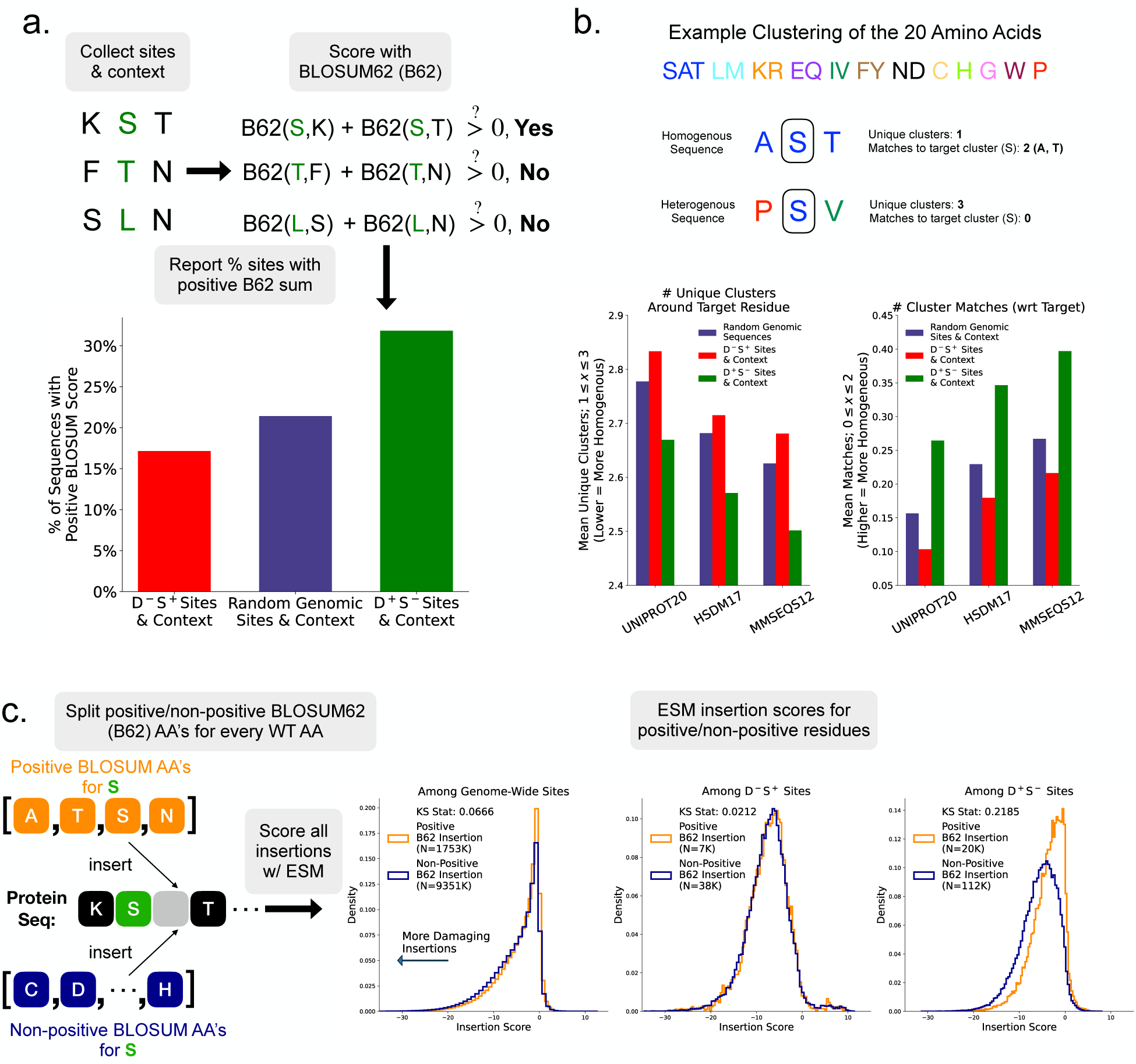
Homogenous sequence contexts mediate differential tolerance to deletions versus substitutions. **(a)** BLOSUM62 (B62) scores for sequence homogeneity. The similarity of a residue and its neighbors is scored with B62. The sign of the sum of B62 scores is used as a proxy of sequence homogeneity. Sequence homogeneity score shown for D^—^S^+^ sites, random sites from the proteome, and D^+^S^—^ sites. **(b)** Example amino acid clustering and scoring of a site’s context. Results for sites (and context) randomly selected from the genome and among sites with differential tolerance. The x-axis ticks are the names of different amino acid clusters^27^ with the number referring to the number of clusters. **(c)** Insertion scan dataset is split between insertions that are similar to its wildtype neighbor (positive B62 score, orange distribution) and all other residues (non-positive B62 score, navy distribution). Insertion scores are reported within these two cases for the full insertion dataset and among sites in the insertion dataset that display differential tolerance.

As an alternative approach, we used amino acid clusters based on shared properties^27–29^ to measure the homogeneity of a given sequence. Consistent with our prior result, we find that the sequence context around D^+^S^—^ sites is more homogenous relative to the contexts of both D^—^S^+^ sites and random background (**Fig. 4b**). This finding is highly robust to the choice of amino acid clustering, the amino acid distribution of the background set, and the length of the sequence context (**Supplementary Figs. S9-11**).

Our analysis suggests that D^+^S^—^ sites are embedded in regions that are robust to small perturbations in length (i.e., deletions) but not perturbations in context. With this understanding, we also anticipated that these regions should be insertion-tolerant, especially when the inserted amino acid is concordant with the local sequence context. For amino acids that are discordant with the sequence context, we expected a more deleterious distribution of effects.

To explore this claim, we returned to our insertion scanning dataset. At every site, the effect of inserting any amino acid was evaluated with ESM. To identify insertions that are concordant with the sequence context, we again used BLOSUM scores: a positive BLOSUM score between the inserted residue and its N-terminal neighbor denotes a concordant insertion, while a non-positive BLOSUM score denotes a discordant insertion. In alignment with our expectations, we observe that D^+^S^—^ sites display differential tolerance to insertions. This differential tolerance is mediated by the concordance (or lack thereof) between the inserted residue and the sequence context (**Fig. 4c**).

In summary, we find homogenous sequence context, as measured by BLOSUM substitution scores and amino acid clusters, to be a distinctive feature of sites that can tolerate deletions but not substitutions (D^+^S^—^). These regions are robust to small perturbations in length but not perturbations in amino acid properties. Our insertion scan data further demonstrates this point, as D^+^S^—^sites are in regions that harbor differential tolerance to insertions, mediated by the identity of the inserted residue. This property is not observed across the full insertion dataset nor among D^—^S^+^ sites.

### Differential tolerance can impact the interpretation of DMS experiments and gnomAD variants

Of the 42,161 human proteins considered in our *in-silico* analysis, 34,504 (nearly 82%) contain at least one site that meets our criteria for differential tolerance. Thus, for the vast majority of human proteins, considering substitutions independently of deletions may generate misleading conclusions on certain sites’ overall mutational tolerance.

As a consequence of this finding, and a strong bias towards substitutions in existing datasets, we hypothesized that results from deep mutational scanning and population genetics could benefit from the joint consideration of deletions alongside substitutions. To substantiate this, we first re-analyzed changes in stability from substitutions and deletions from Tsuboyama et al. (2023), where we identified many instances of differential tolerance within the experiment (**Fig. 2b**). We observed certain domains, such as the C domain of the staphylococcal protein A (PDB accession 6SOW), as strongly overrepresented among D^—^S^+^ sites. Consistent with our understanding of this property, the C domain is almost exclusively composed of alpha-helices; 44 of the domain’s 56 residues are mapped to one of three helices.

In this domain, the glycine at position 23 meets our criteria for D^—^S^+^ differential tolerance. Particularly, this position is more substitution-tolerant than over 96% of sites measured in Tsuboyama et al. (2023). On the other hand, the site is more deletion-intolerant than over 90% of sites in the experiment. Computational structure prediction with ESMFold^11^ helps to clarify the site’s differential tolerance. While the wildtype structure holds all helices at approximately the same orientation, the *Δ*G23 mutant modifies the positioning of the third helix (**Fig. 5a**). Moreover, the deletion decreases overall model confidence, a pattern observed across all D^—^S^+^ sites in the study (**Fig. 5b**). Because the C domain mediates contact with human antibodies,^30,31^ this deletion could modify the immune system response to the staphylococcus pathogen. Such a result would not be possible to generate from substitutions alone, however, given the site’s robustness to substitutions.

**Figure 5:**
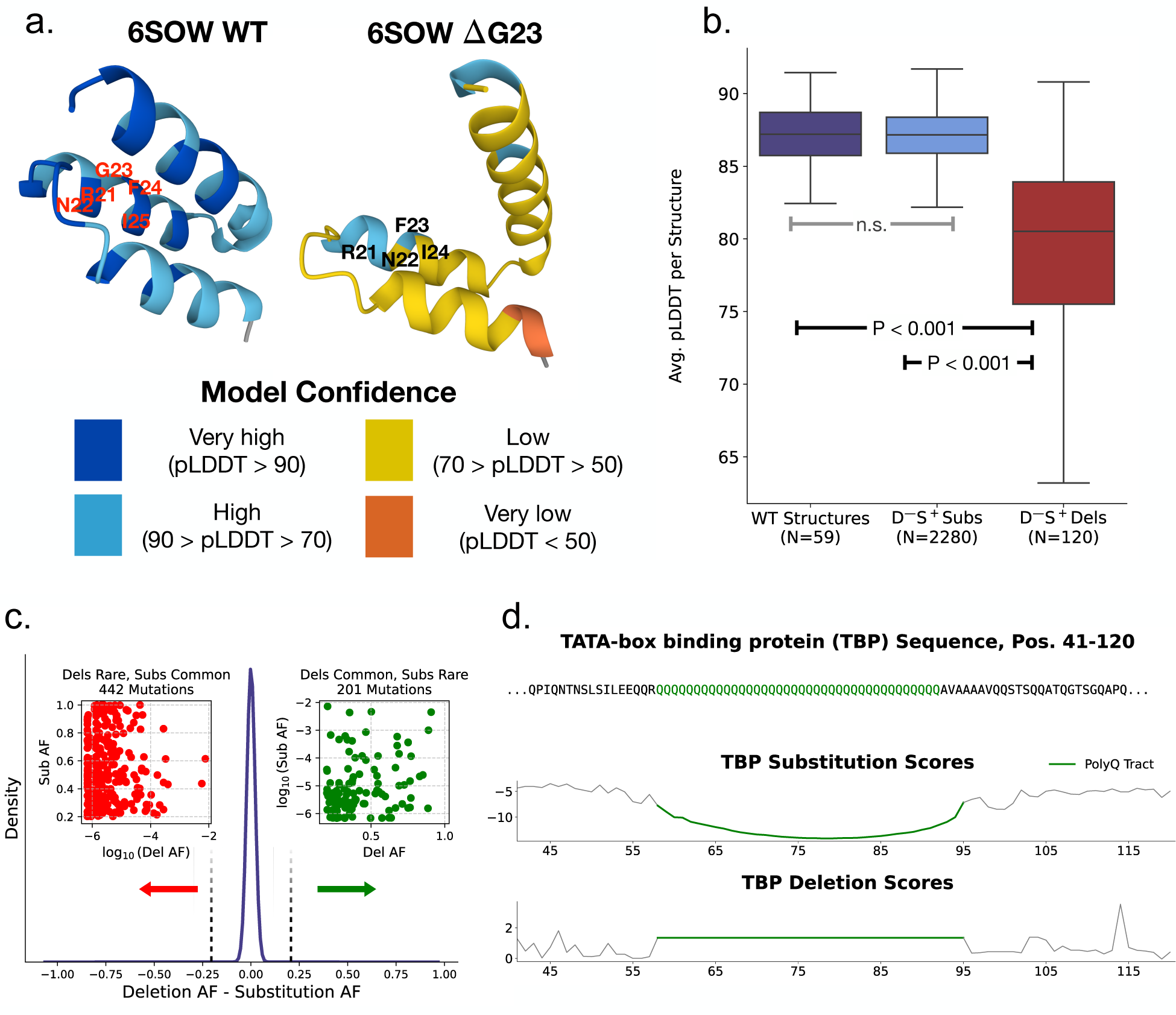
Differential tolerance is observed in mutational scanning and population genetics data. **(a)** 6SOW wildtype structure (left) and *Δ*G23 (right), colored by pLDDT (bottom). **(b)** ESMFold pLDDT for all Tsuboyama et al. WT structures containing a D^—^S^+^ site, for all possible substitutions among D^—^S^+^ sites, and for all deletions among D^—^S^+^ sites. P-values reported from t-test. **(c)** Difference in gnomAD allele frequency between substitutions and deletions occurring at the same position. **(d)** TATA-box binding protein sequence (top), ESM-predicted substitution scores by position (middle), and ESM-predicted deletion scores by position (bottom). Deletion & substitution scores close to zero are considered neutral, more negative scores are considered more damaging.

It is also possible to identify differences between substitution and deletion effects in population genetics data, where allele frequency can proxy mutational tolerance^32^. In particular, we identified 303,603 positions in gnomAD where both a missense variant and an in-frame deletion are observed. While the difference in allele frequency between a substitution and a deletion tends to be very small, we also observed many instances where one mutation is rare (AF < 1%) and the other is very common (AF > 20%, **Fig. 5c**). For example, the transcription factor TATA-box binding protein has a common (allele frequency >0.4 in gnomAD) in-frame deletion of a glutamine at position 95.

However, the substitution Q95H is extremely rare, with an allele frequency of 0.000002. This glutamine is located within a 38-residue polyglutamine tract (**Fig. 5d**), in agreement with our result that deletion-tolerant, substitution-intolerant sites tend to be embedded within homogenous sequence contexts. The Q95 deletion is annotated as benign in ClinVar, while Q95H is unannotated^33,34^. However, despite these mutations occurring at the same site, they may not have the same clinical impact; applying ESM-1b to the TBP sequence reveals that the polyglutamine tract, including Q95, is generally substitution-intolerant and deletion-tolerant (**Fig. 5d**).

Overall, these results underscore the value of considering substitutions alongside deletions in studies of protein fitness. Evidence of differential tolerance can be found in existing experimental and population genetics datasets. If deletion effects were excluded from these analyses, an incomplete understanding of mutational tolerance would be drawn for sites like 6SOW G23 and TBP Q95.

## Discussion

Deletions are a critical class of mutations in protein evolution, engineering, and human genetics. Despite this, their genome-wide effects remain poorly understood, particularly relative to substitutions. To further our understanding of the relationship between deletions and substitutions, we used the ESM-1b protein language model to predict the effects of all amino acid deletions in the human proteome.

We used ESM-1b to compare deletion versus substitution tolerance genome-wide. The scale of this analysis is unprecedented in the literature and allowed us to identify hundreds of thousands of sites that display differential tolerance. The existence of these sites suggests that substitution effects, while correlated with deletion effects in aggregate, are insufficient to fully capture the tendencies of deletion intolerance.

Among sites that are deletion-intolerant and substitution-tolerant, we find secondary structural elements to be an important predictive feature. In particular, sites contained within alpha-helices are strongly overrepresented among D^—^S^+^ sites relative to their background frequency in the human proteome. On the other hand, sites that are deletion-tolerant but substitution-intolerant tend to be found within more homogenous sequence contexts than D^—^S^+^ sites and random sites from the proteome. Our results are the first to provide genome-wide evidence for differential tolerance between substitutions and deletions across *in-silico*, experimental, and population datasets, and establish protein language models as a powerful tool for biological hypothesis generation and analysis.

Our results pose numerous areas for further exploration. To begin, we identify certain features associated with differential tolerance, namely alpha-helices and homogenous sequence context. However, not every site that displays differential tolerance possesses these features. Relatedly, not every site with these features displays differential tolerance. Resolving these ambiguities will create a more nuanced view of why differential tolerance does (and does not) occur.

As demonstrated, the principle of differential tolerance can help to further refine clinical annotations. In particular, existing clinical guidelines draw an explicit link between mutations occurring at the same site in a protein^35^. While refining such guidelines is beyond the scope of this study, we argue that differential tolerance may challenge the direct transfer of clinical annotations between a substitution and a deletion at the same site.

Lastly, our results demonstrate the power of protein language models to proxy an experimental system. This class of models has already demonstrated highly accurate predictive performance across relevant biological domains. However, we demonstrate their utility in answering second-order biological questions, such as the comparison between different types of mutations. Our analysis showcases an exciting potential for these tools to supplement limited experimental data to generate new biological hypotheses.

## Methods

### Accessing ESM

ESM-1b was accessed pretrained from GitHub - facebookresearch/esm: Evolutionary Scale Modeling (esm): Pretrained language models for proteins using the esm.pretrained.esm1b_t33_650M_UR50S module.

### Scoring substitutions with ESM

We scored substitutions with ESM via the log-likelihood ratio (LLR), which compares the log-likelihood of the mutant residue against the log-likelihood of the reference residue. Code for scoring substitutions via the LLR can be found at https://github.com/ntranoslab/esm-variants/blob/main/esm_score_missense_mutations.py. Because each site has 19 substitution scores but only one deletion score, we aggregated substitution effects at the site level to compare the measurements one-to-one. Among the different aggregation functions tested, we found that a site’s average LLR most accurately predicts ClinVar annotations compared to the site’s median, max, and min LLR. As such, we chose the average LLR to represent a site’s substitution tolerance.

### PLLR framework for scoring mutations with ESM

The pseudo log-likelihood ratio (PLLR) is defined as the ratio of the pseudo-log-likelihoods of the *entire* mutant sequence and the *entire* wildtype sequence, as estimated by ESM^3^. All deletions and insertions in this study were scored with the PLLR. We also used the PLLR metric for substitutions in **Fig. 3c** and **Supplementary Fig. S4.** Code for scoring mutations via the PLLR can be found at https://github.com/ntranoslab/esm-variants/blob/main/esm_score_multi_residue_mutations.py.

### Scoring long sequences with ESM

To overcome ESM-1b’s hard-coded 1,022 sequence length limit, we re-employed our existing tiling framework^3^.

### Visualizing the distribution of deletion effects

Prior research indicates that the absolute PLLR (aPLLR) is the most accurate predictor of benign/pathogenic clinical annotations^3^. As such, we conducted all deletion analyses in Fig. 3, including the data partitioning, using the aPLLR. To aid with visualization, however, we *plot* the aPLLR’s on the -log10 scale in **Fig. 2a** and **Fig. 5d**. This facilitates the visual inspection of deletion effects across the aPLLR range.

### Accessing ClinVar annotations for substitutions and deletions

The median ESM LLR of ClinVar benign and ClinVar pathogenic substitutions were taken from https://github.com/ntranoslab/esm-variants/blob/main/benchmarks/ClinVar_gnomAD_benchmark_with_predictions.csv. For deletions, we used https://github.com/ntranoslab/esm-variants/blob/main/benchmarks/ClinVar_indel_benchmark_with_predictions.csv and filtered to only deletions (removing insertions and del-ins).

### Secondary structural element prediction

All AlphaFold human .pdb predictions were extracted from https://alphafold.ebi.ac.uk/download. Secondary structural elements were predicted using the Bio.PDB.DSSP module from the BioPython library (https://biopython.org/docs/1.75/api/Bio.PDB.DSSP.html). We grouped all sites with an S, H, or I annotation into Helices, B and E into Strands, and S and T into Loops.

### Insertion and substitution scanning dataset

We selected 1,632 proteins whose structures were approximately 50% alpha-helix, as estimated by DSSP. All proteins selected correspond to canonical transcripts from the AlphaFold database. For each site, we used the PLLR metric to estimate the effect of both (a) inserting a mutant residue to the right of the wildtype residue, and (b) substituting the wildtype residue to the mutant residue. In **Fig. 3c**, we excluded synonymous substitutions and insertions of the same amino acid as wildtype to avoid biasing the results towards substitutions being more damaging.

### Analysis of experimental stability measurements

DMS data was downloaded from Tsuboyama et al. (https://zenodo.org/records/7992926). The average substitution **!!**G was computed per position. Variants not mapping to a PDB identifier or absent of any substitution, insertion, or deletion ΔΔG values were removed.

- ● DMS ΔΔG data:

processed_K50_dG_datasets/Tsuboyama2023_Dataset2_Dataset3_20230416.csv

- ● Secondary structure data: Data_tables_for_figs/dG_site_feature_Fig3.csv

### Accessing gnomAD data

All gnomAD v4 exome variants were downloaded from https://gnomad.broadinstitute.org/downloads, and filtered down to any missense mutations or single in-frame amino acid deletions. Additional filtering was applied to remove any variants with an allele frequency of 0. To find sites containing both a deletion and substitution(s), we merged the deletion data and substitution data on an Ensembl Protein ID (ENSP*) and a position. UniProt ID’s were added via gProfiler convert https://biit.cs.ut.ee/gprofiler/convert.

## Data availability statement

To aid with reproducibility and downstream research, we released all supplementary data and code for public access at https://github.com/ntranoslab/diff-tol. Data on genome-wide differential tolerance can be downloaded and explored at https://huggingface.co/spaces/goldmangrant/diff-tol.

## Supplementary Figures

**Supplementary Figure S1:**
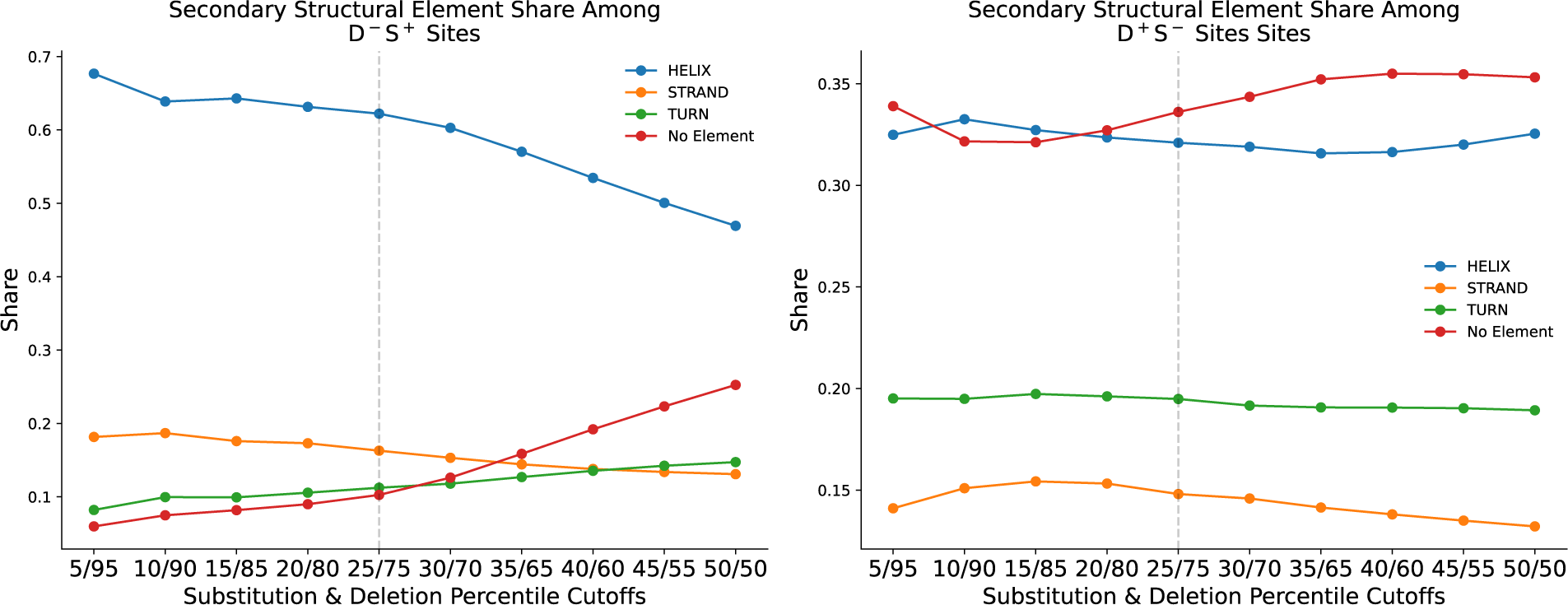
Secondary structural element shares in D^—^S^+^/ D^+^S^—^ quadrants as a function of percentile cutoff. Share of sites belonging to each DSSP category in the deletion-intolerant, substitution tolerant (D^—^S^+^) quadrant or deletion tolerant, substitution-intolerant (D^+^S^—^) quadrant when varying the percentile cutoff. The 25th/75th percentile cutoff was selected for the main analysis.

**Supplementary Figure S2:**
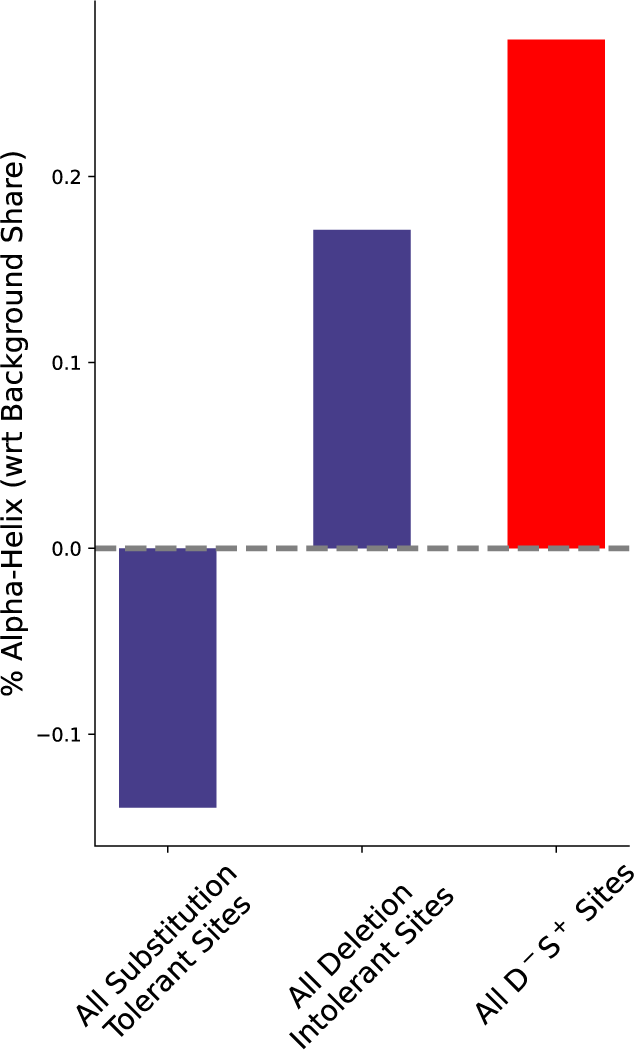
Share alpha-helix within different data partitions. Relative to the background frequency in the data, the share of alpha-helices among all substitution-tolerant sites, all deletion-intolerant sites, and all D^—^S^+^ sites.

**Supplementary Figure S3:**
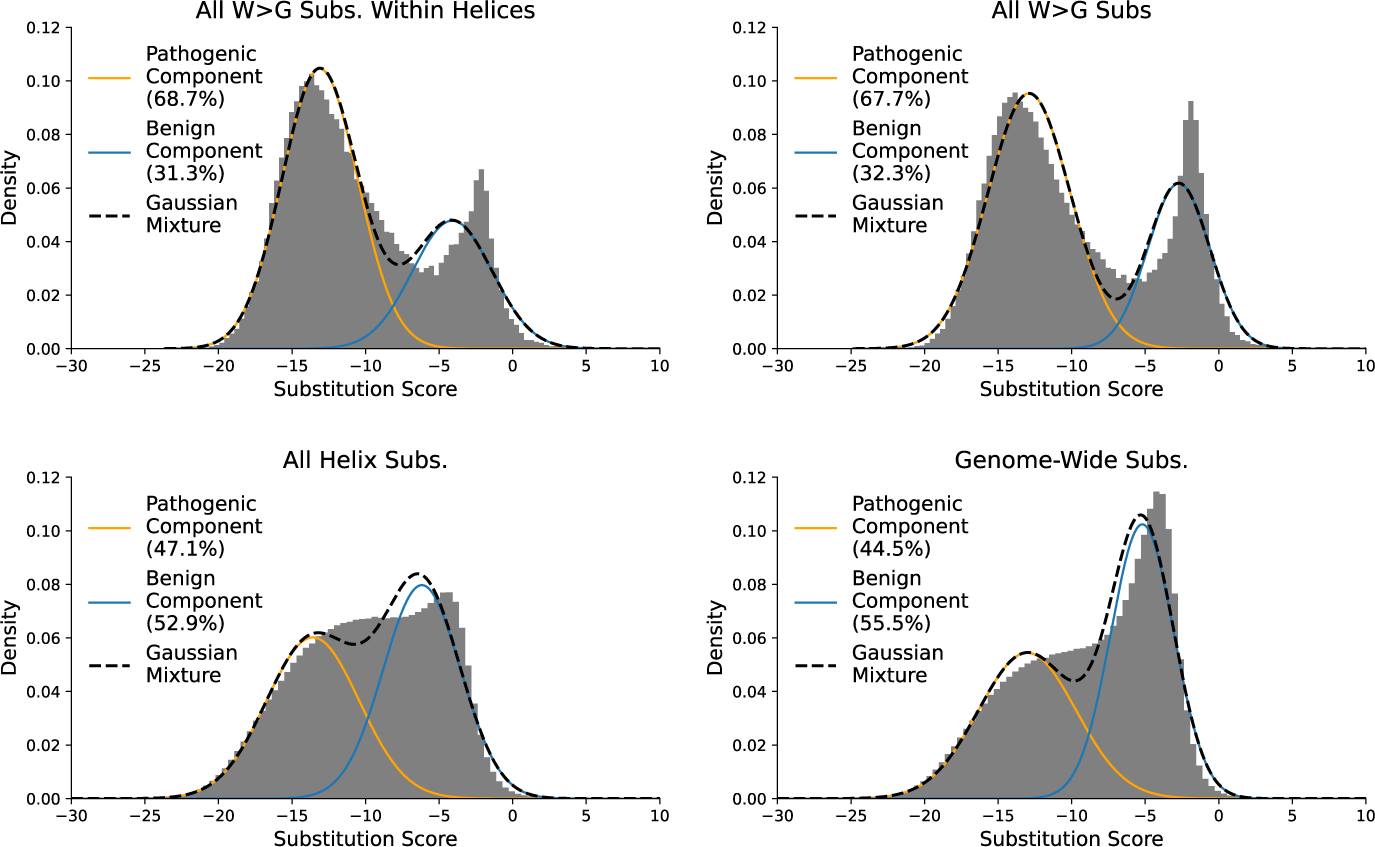
Decomposing W>G substitutions into pathogenic and benign components. Probability density of substitutions scores for W>G substitutions within helices (top left), all W>G substitutions (top right), all helix substitutions (bottom left), and among genome-wide substitutions (bottom right). For each dataset, a two-component Gaussian mixture is fit to the data, and the weights of the mixture are reported.

**Supplementary Figure S4:**
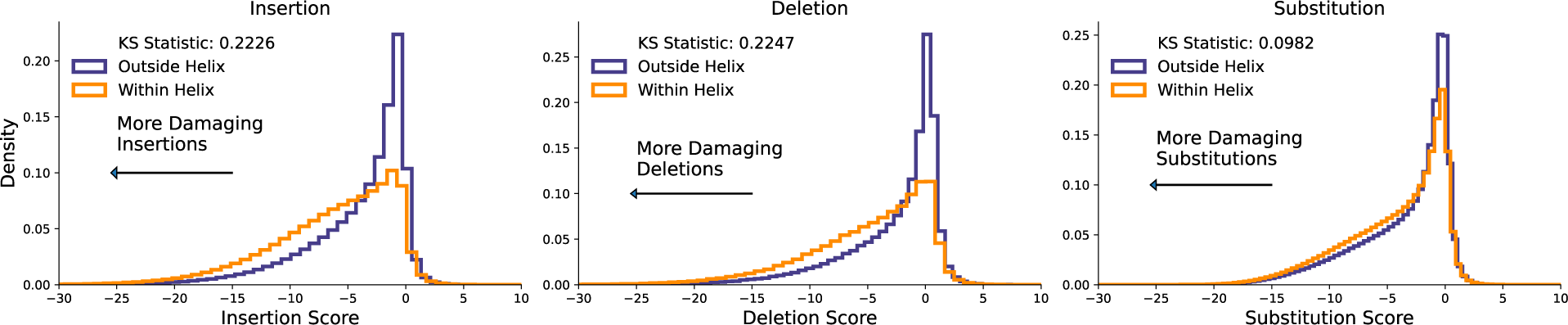
Tolerance to insertions, deletions, and substitutions within versus outside helices. Insertions, deletions, and substitutions were scored with the ESM PLLR metric for 1,632 proteins. The effect of the mutation within versus outside a helix is shown.

**Supplementary Figure S6:**
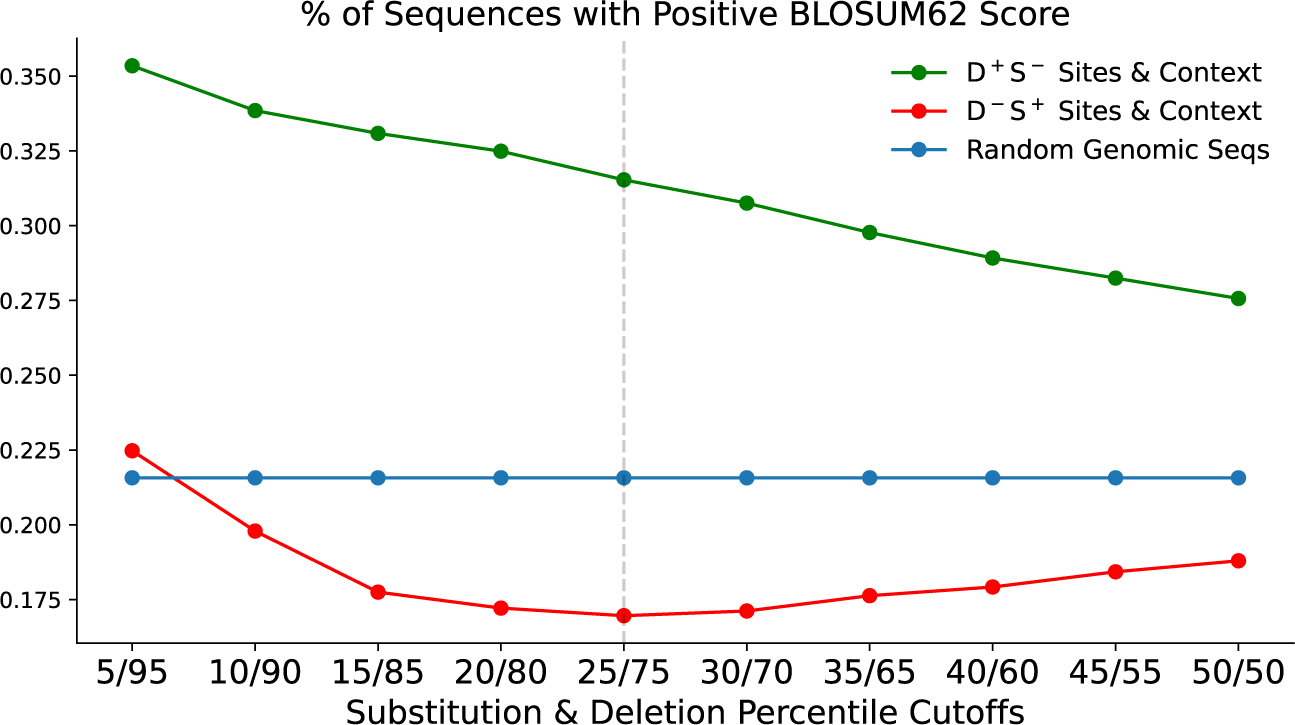
Percent of sequences with positive BLOSUM62 (B62) scores as a function of percentile cutoff. Approximating sequence homogeneity with B62 scores among random genomic sequences and within each quadrant when varying the percentile cutoff used to construct the quadrants.

**Supplementary Figure S7:**
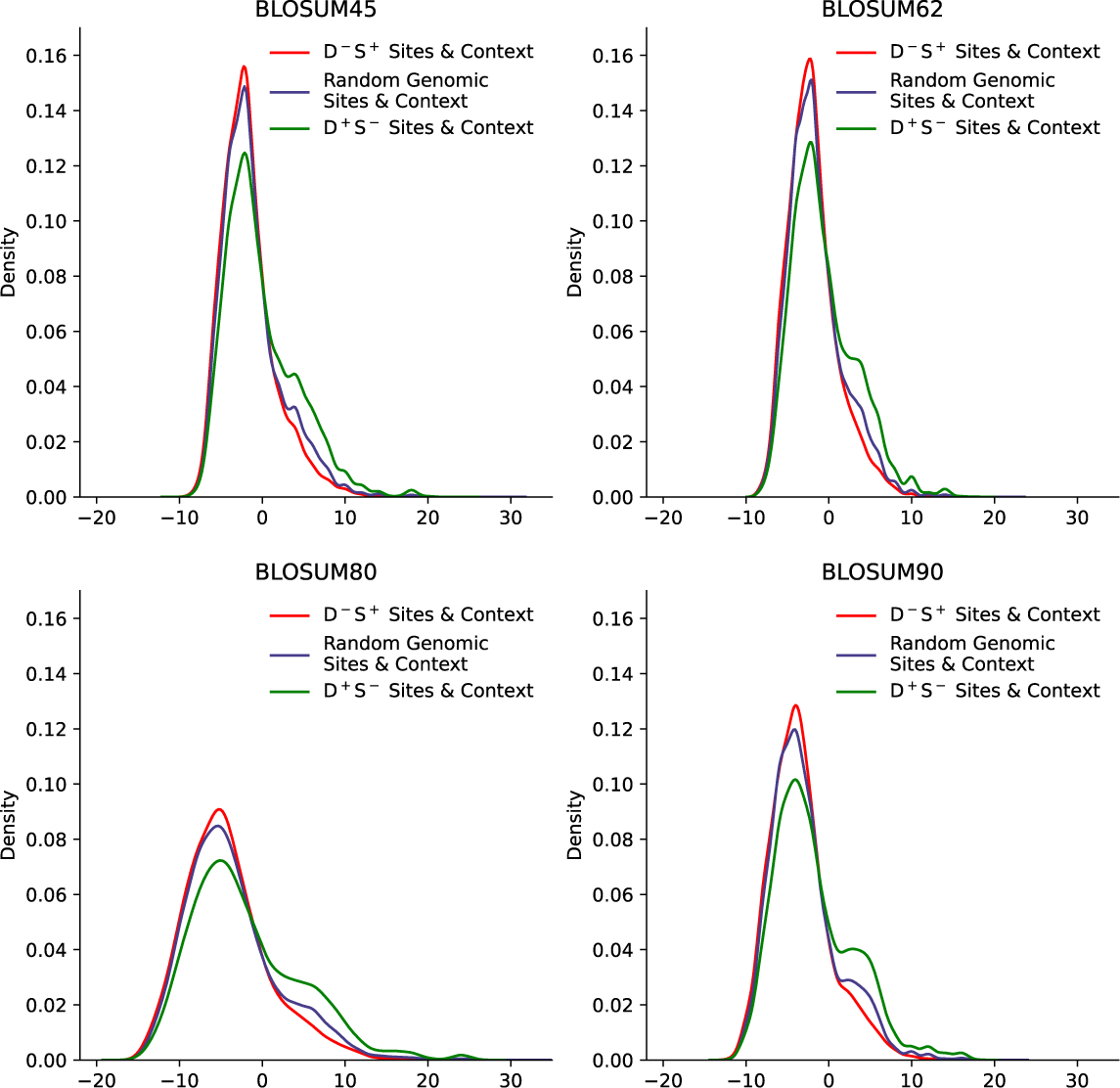
Distribution of BLOSUM sequence sums for multiple substitution matrices. Scoring sequence homogeneity using BLOSUM substitution matrices. A positive value indicates a more homogenous sequence context.

**Supplementary Figure S8:**
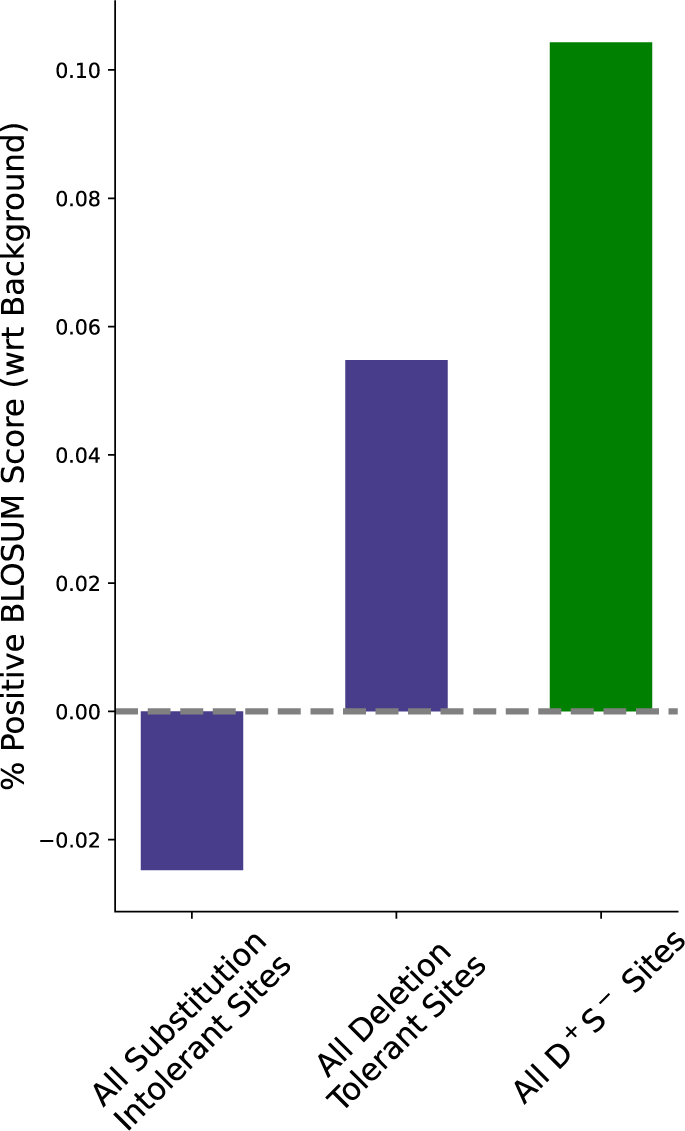
Percent positive BLOSUM score within different data partitions. Relative to the background share in the data, the percent of sites and context with a positive BLOSUM score among all substitution-intolerant sites, all deletion-tolerant sites, and all D^+^S^—^ sites.

**Supplementary Figure S9:**
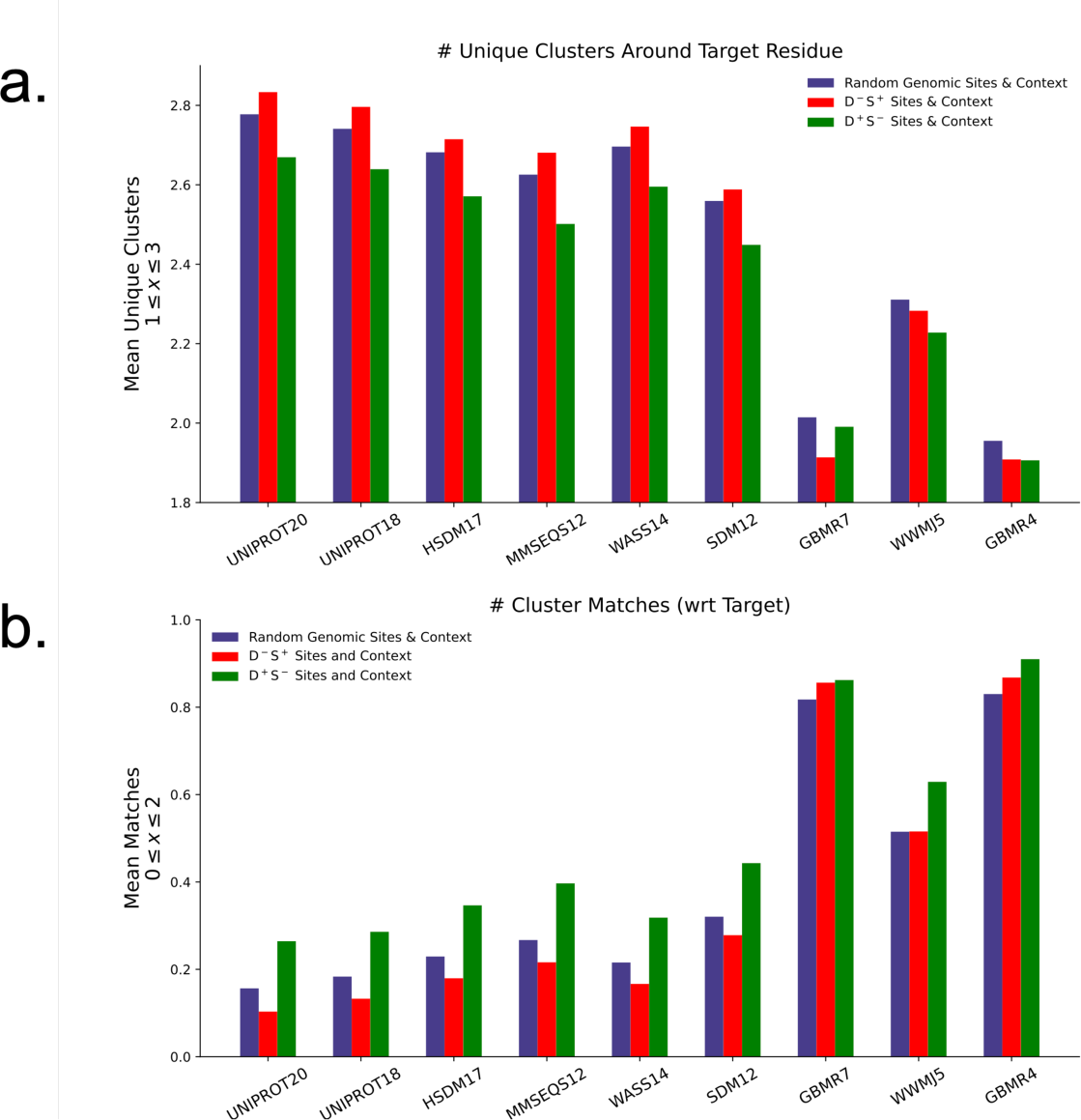
Scoring sequence homogeneity across an expanded set of amino acid clusterings. **(a)** Counting the number of unique clusters represented in a sequence window among genome wide sites (blue bars) and all sites in the D^—^S^+^ and D^+^S^—^ quadrant (red and green bars, respectively) for an expanded set of amino acid clustering approaches. The number of clusters is given in the cluster name. **(b)** Counting the number of cluster matches to the target residue in a sequence window among genome wide sites (blue bars) and all sites in the D^—^S^+^ and D^+^S^—^ quadrant (red and green bars, respectively) for an expanded set of amino acid clustering approaches. The number of clusters is given in the cluster name.

**Supplementary Figure S10:**
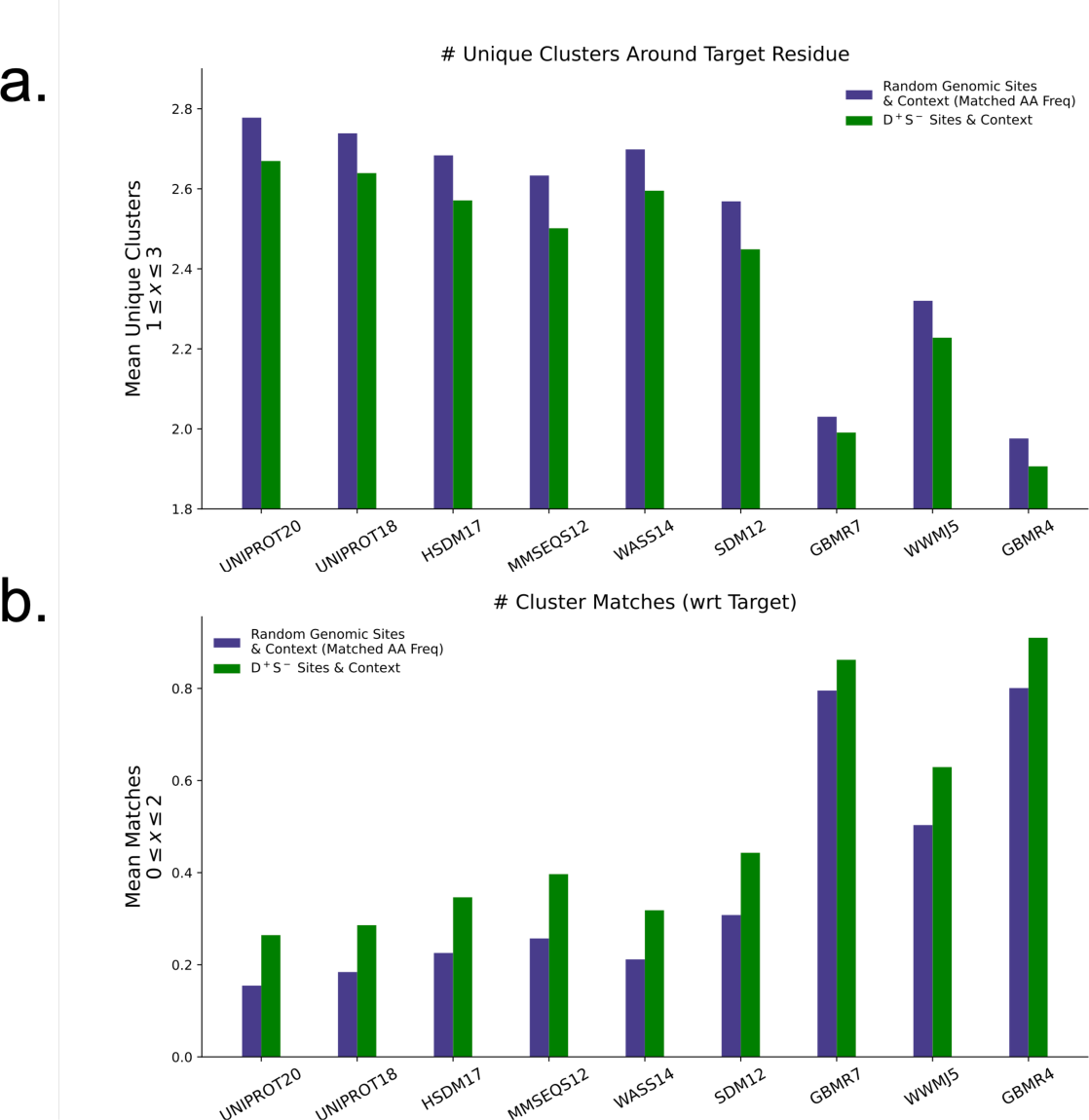
Scoring sequence homogeneity with a dataset matched on amino acid frequency. Sequence homogeneity scores between sites in the D^+^S^—^ quadrant versus a genome-wide set. The genome-wide set was redrawn to match the frequency of amino acids in the D^+^S^—^ quadrant. This adjusts for the tendency of certain residues (i.e., glycine, proline, tryptophan) to be in their own cluster across alphabets, which makes sequences that contain them appear more heterogeneous.

**Supplementary Figure S11:**
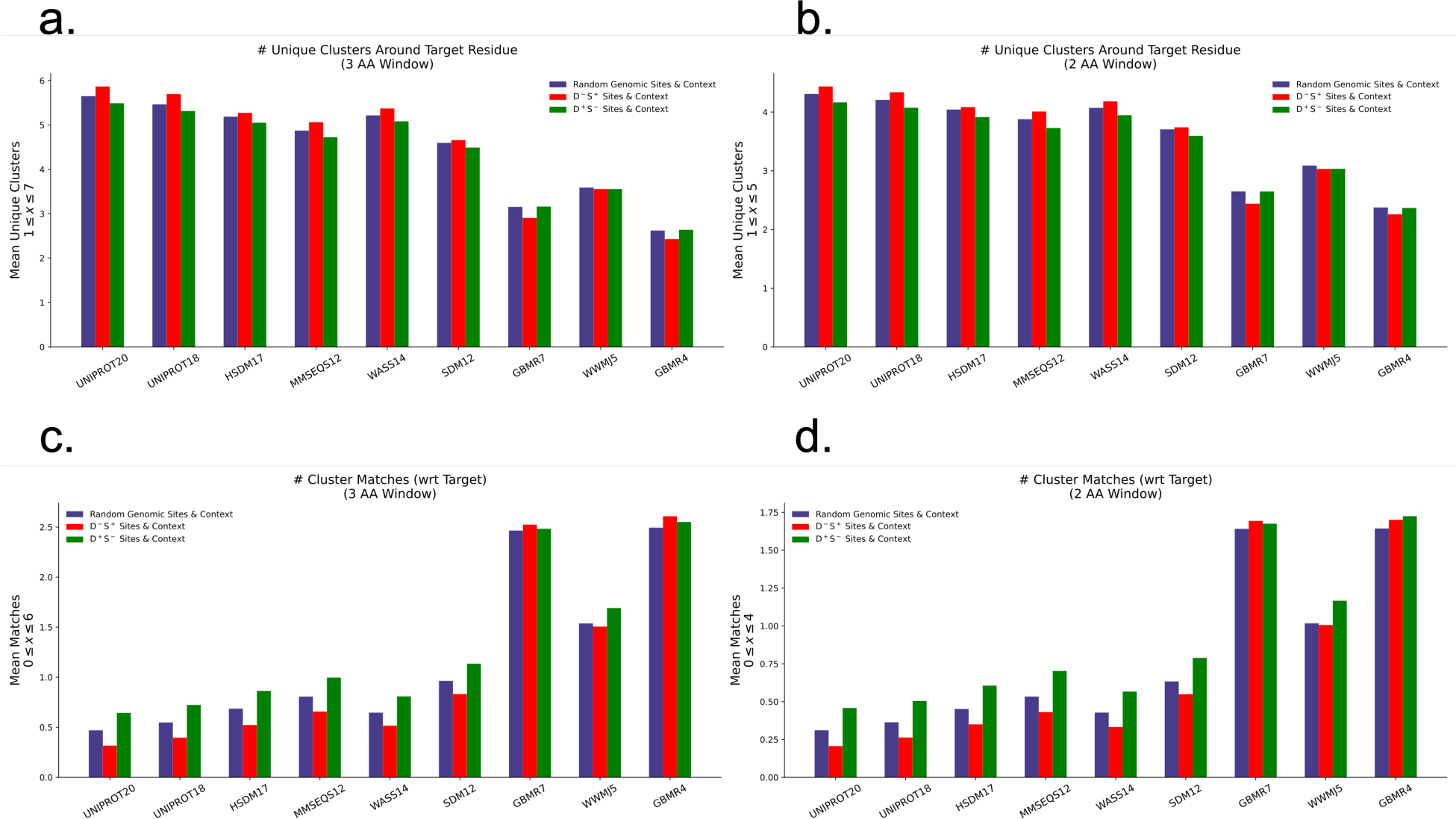
Scoring sequence homogeneity across a wider sequence range. **(a)** Number of unique clusters represented around a three amino acid window (in either direction) of the target residue. **(b)** Number of unique clusters represented around a two amino acid window (in either direction) of the target residue. **(b)** Matches to target residue around a three amino acid window (in either direction) of the target residue. **(d)** Matches to target residue around a two amino acid window (in either direction) of the target residue.

